# Predicting Fungal Contaminants for Space Missions Using Proteome-Wide Screening for Protein Orthologs

**DOI:** 10.64898/2026.08.22.746478

**Authors:** Ashish Mahabal, Vannsh Jani, S. George Djorgovski, Nitin K. Singh, Swati Bijlani

**Author notes:** **Preprint of an accepted manuscript.** This article has been accepted for publication in *Astrobiology*. This is the author’s accepted manuscript; it has not been copyedited or typeset by the publisher. The version of record will be available from the journal, and this posting will be updated with a link to it on publication.

## Abstract

Fungal contamination poses a growing threat to spacecraft integrity, crew health, and planetary protection efforts. We describe a scalable and interpretable pipeline for identifying fungi with adaptation potential to spaceflight-associated stress conditions such as extreme temperatures, radiation levels, etc., and pathogenicity risks. Starting with proteins known to confer stress resistance, we identify orthologs across over fifteen hundred fungal species and evaluate their contamination potential via comparative proteome analysis. Our pipeline integrates proteins with known functional inference, cross-database proteome matching, and identity-based scoring to generate a ranked list of fungal species of concern. We apply this approach to detections from spacecraft assembly facilities, highlighting species with combined stress-tolerance and pathogenic potential. This study establishes a foundation for future AI-based risk assessments that can scale to orders of magnitude more fungal species, thus laying the foundation for systematic identification and assessment of fungal contaminants with potential adaptation and pathogenicity risks in spaceflight environments, thereby supporting contamination control strategies for future space missions. We also present an interactive visual online tool for researchers to trivially check the contamination potential of species in their own samples.

## 1 Introduction

Fungi are increasingly recognized as critical yet under-monitored agents of contamination in both terrestrial and spaceflight environments (dos Santos et al., 2025). Reports of fungal infections and material degradation span hospitals, laboratories, and even spacecraft assembly facilities (Camacho et al., 2009; Cock et al., 2009; De Middeleer et al., 2019; Bijlani et al., 2022; Fisher et al., 2022; Xu, 2022; Fisher and Denning, 2023; Lass-Flörl and Steixner, 2023; Fisher et al., 2024; Konkel Neabore, 2024; Seidel et al., 2024; Benardini and Lalime, 2025; Kadariswantiningsih et al., 2025). Even tightly controlled cleanrooms and the International Space Station (ISS) have shown persistent fungal presence (Singh et al., 2018; Zhang et al., 2018; De Middeleer et al., 2019), raising concern for long-duration missions where closed habitats, radiation, and limited resources may amplify microbial risks.

This is true despite NASA and other space agencies having strict mission-specific standards for determining possible microbial contamination (Benardini and Lalime, 2025), raising concern for future deep space missions. Fungal species possess exceptional survival traits such as resistance to desiccation, radiation extremes, temperature extremes, and nutrient scarcity that make them uniquely suited to endure spaceflight-associated stressors (Bijlani et al., 2021). These same mechanisms can compromise spacecraft materials, alter habitat microbiomes, and threaten astronaut health. However, comprehensive fungal risk evaluation remains difficult because of incomplete proteome annotations, uneven metadata quality, and sparse experimental characterization across the fungal tree of life (Simpson et al., 2021).

To address this gap, we developed a systematic, proteome-wide screening pipeline that infers contamination potential using orthologs of well-characterized stress and virulence-associated proteins. Starting from known fungal proteins linked to antimicrobial resistance, biofilm formation, pathogenicity, radiation tolerance, spore formation, and thermophilicity, we identify homologous sequences across more than fifteen hundred fungal species. This approach enables scalable inference of functional capabilities even in newly sequenced or poorly annotated organisms.

By quantifying sequence similarity and combining data from multiple public repositories e.g. NIH ^1^, MycoCosm (Grigoriev et al., 2014), UniProt ^2^, GOLD (Mukherjee et al., 2025), and NCBI RefSeq, our workflow establishes an extensible foundation for AI-based contamination risk modeling. Such models could integrate environmental context, abundance, and stress data to anticipate microbial threats under spaceflight and planetary protection scenarios.

For this pilot study we have created a curated list of over fifteen hundred fungal species. We have incorporated the list of organisms and their traits on a web application that allows users to identify fungal species from our curated list that potentially survive extreme conditions like extreme temperatures, high radiation levels, etc. Users input datasets containing numbers of reads for various organisms at one or more locations, and are returned the contamination potential for each location, based on thresholds that the user can decide, pointing out organisms and their potentially harmful properties. This work lays the foundation for systematic identification and assessment of fungal contaminants with potential adaptation and pathogenicity risks in spaceflight environments, thereby supporting contamination control strategies for future missions by NASA as well as other space agencies. For instance, one such NASA requirement (NID 8715.129, 1.3.1 - 1.3.6) is to identify, develop, and implement in-flight microbial monitoring and diagnostic tools to support research and crew health during Gateway, Lunar, and Mars missions ^3^. Another is connected with planetary protection (PP) – to ensure the prevention of potentially harmful consequences for humans and the Earth’s environment following the return of extraterrestrial samples from restricted designations (NASA-STD-8719.27).

## 2 Materials and Methods

This section describes the workflow developed to identify fungal species with potential for spaceflight-associated stress adaptation and pathogenicity. The process includes selection of representative stress-associated proteins, compilation of fungal proteomes from multiple repositories, ortholog identification through BLAST-based alignments, and derivation of contamination potential scores. Each step was designed to be scalable and reproducible for integration into future AI-driven risk assessment frameworks.

### 2.1 Selection of Stress-Associated Proteins

We identified seven traits fungi exhibit that can be potentially harmful under space conditions as documented in various NASA studies - see Table 1. We then identified well-studied fungal species that demonstrate these traits, and then identified proteins that are associated with those traits in the selected species (See Table 2). The phyla for the selected species are given in Table 3. For the 36 proteins thus identified, we looked for FASTA sequences in the UniProtKB portal and found 25 there. We restrict ourselves to these 25 proteins across six species and six traits in this study. We do not claim that the species or proteins we have selected are fully representative of the traits under consideration. Our methodology is scalable and extensible so that it will not be difficult to extend to more organisms and more proteins in the future (by looking at, for instance, the NCBI database, which reveals FASTA sequences for five additional proteins from our initial selection of 36). With future scalability in mind, we identify many protein orthologs that can be used as bonafide proteins to extend the set of organisms with the properties we are interested in. The second and third parts of Table 2 list proteins we have excluded from ortholog searches in this study. This modular design ensures that the workflow can be expanded easily to incorporate new traits and proteins as annotations improve, and more datasources are incorporated.

**Table 1:** NASA traits and their provenance. Throughout this work, for the traits/properties given in the first column, we will use the shortforms given in the second column.

| Trait | Shortform | NASA Document | Year |
| --- | --- | --- | --- |
| Antimicrobial resistance | AMR | Microbiology in Spaceflight Overview<sup>a</sup> | 2025 |
| Biofilm formation | BF | Microbiology in Spaceflight Overview<sup>a</sup> | 2025 |
| Human pathogenicity | HP | Microbiology in Spaceflight Overview<sup>a</sup> | 2025 |
| Psychrophily | PS | Microbial Examination of Space Hardware<sup>b</sup> | 2019 |
| Radiation resistance | RAD | Advancing Shotgun Metagenomics<sup>c</sup> | 2024 |
| Spore formation | SF | Planetary Protection Handbook<sup>d</sup> | 2024/2025 |
| Thermophilicity | TH | Microbial Examination of Space Hardware<sup>b</sup> | 2019 |

<sup>a</sup> https://www.nasa.gov/wp-content/uploads/2025/05/ochmo-tb-046-microbiology-rev-a.pdf
<sup>b</sup> https://explorers.larc.nasa.gov/2019APSMEX/SMEX/pdf_files/NASA-HDBK-6022b.pdf
<sup>c</sup> https://sma.nasa.gov/docs/default-source/sma-disciplines-and-programs/planetary-protection/kasthuri-venkateswaran_venkat-metagenome-workshop-at-ames-2024-presentation-nov-19-2024.pdf
<sup>d</sup> https://ntrs.nasa.gov/api/citations/20240016475/downloads/PlanetaryProtection_Hdbk_2024_Final_For508ing_comp.pdf

**Table 2:** Traits, organisms, and proteins selected for this study. The first section lists proteins with FASTA sequences found in UniProt and included in the study (this does not include any psychrophilic protein). The second section lists proteins with FASTA sequences available in NCBI but not considered further in this study. The last section lists proteins for which we could not find FASTA sequences in either UniProt or NCBI, but likely exist elsewhere. This collection demonstrates the scalability of our workflow to include additional proteins across diverse functional categories.

| Property | Organism | Protein |
| --- | --- | --- |
| **FASTA sequences available in UniProt** |  |  |
| Antimicrobial resistance | *Candida albicans* | Cdr1 |
| Antimicrobial resistance | *Candida albicans* | Erg11 |
| Antimicrobial resistance | *Candida albicans* | Mdr1 |
| Antimicrobial resistance | *Candida albicans* | Mrr1 |
| Antimicrobial resistance | *Candida albicans* | Tac1 |
| Antimicrobial resistance | *Candida albicans* | Upc2 |
| Biofilm formation | *Candida albicans* | Als3 |
| Biofilm formation | *Candida albicans* | Bcr1 |
| Biofilm formation | *Candida albicans* | Czf1 |
| Biofilm formation | *Candida albicans* | Efg1 |
| Biofilm formation | *Candida albicans* | Erg251 |
| Biofilm formation | *Candida albicans* | Flo8 |
| Biofilm formation | *Candida albicans* | Hwp1 |
| Biofilm formation | *Candida albicans* | Ndt80 |
| Biofilm formation | *Candida albicans* | Tec1 |
| Human pathogenicity | *Candida albicans* | Rim101 |
| Human pathogenicity | *Candida albicans* | Sap5 |
| Human pathogenicity | *Cryptococcus neoformans* | Lac1 |
| Human pathogenicity | *Cryptococcus neoformans* | Plb1 |
| Radiation resistance | *Saccharomyces cerevisiae* | Rad51 |
| Spore formation | *Coprinopsis cinerea* | srr1 |
| Spore formation | *Emericella nidulans* | AbaA |
| Spore formation | *Emericella nidulans* | BrlA |
| Spore formation | *Emericella nidulans* | WetA |
| Thermophilicity | *Aspergillus fumigatus* | Hsp90 |
| **FASTA sequences available in NCBI** |  |  |
| Antimicrobial resistance | *Candida glabrata* | Fsk1 |
| Antimicrobial resistance | *Candida glabrata* | Pdr1 |
| Psychrophily | *Antarctomyces psychrotrophicus* | Anpibp |
| Psychrophily | *Glaciozyma antarctica* | Afp4 |
| Thermophilicity | *Cryptococcus neoformans* | Tps1 |
| **FASTA sequences not found in UniProt or NCBI** |  |  |
| Human pathogenicity | *Cryptococcus neoformans* | Cap59 |
| Psychrophily | *Cryomyces antarcticus* | Ole1 |
| Radiation resistance | *Exophiala dermatitidis* | Ku70 |
| Radiation resistance | *Exophiala dermatitidis* | Pks1 |
| Thermophilicity | *Aspergillus fumigatus* | HsfA |

**Table 3:**
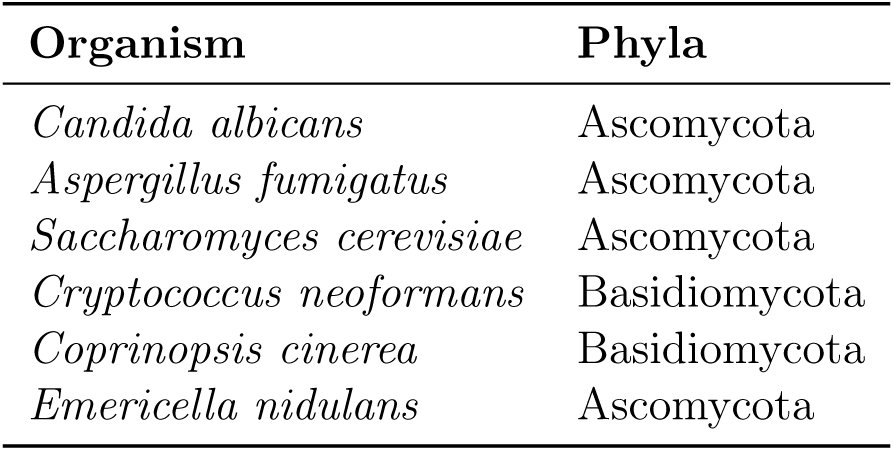
Classification of fungal organisms in which query proteins were found.

### 2.2 Workflow

The overall workflow is summarized in Figure 1. The process involves: (1) compilation of fungal species, (2) construction of organism-specific BLAST databases from retrieved proteomes, and alignment of query proteins against these databases using BLASTp to identify putative orthologous sequences and (3) compute identity scores. Each stage is described in detail below.

**Figure 1:**
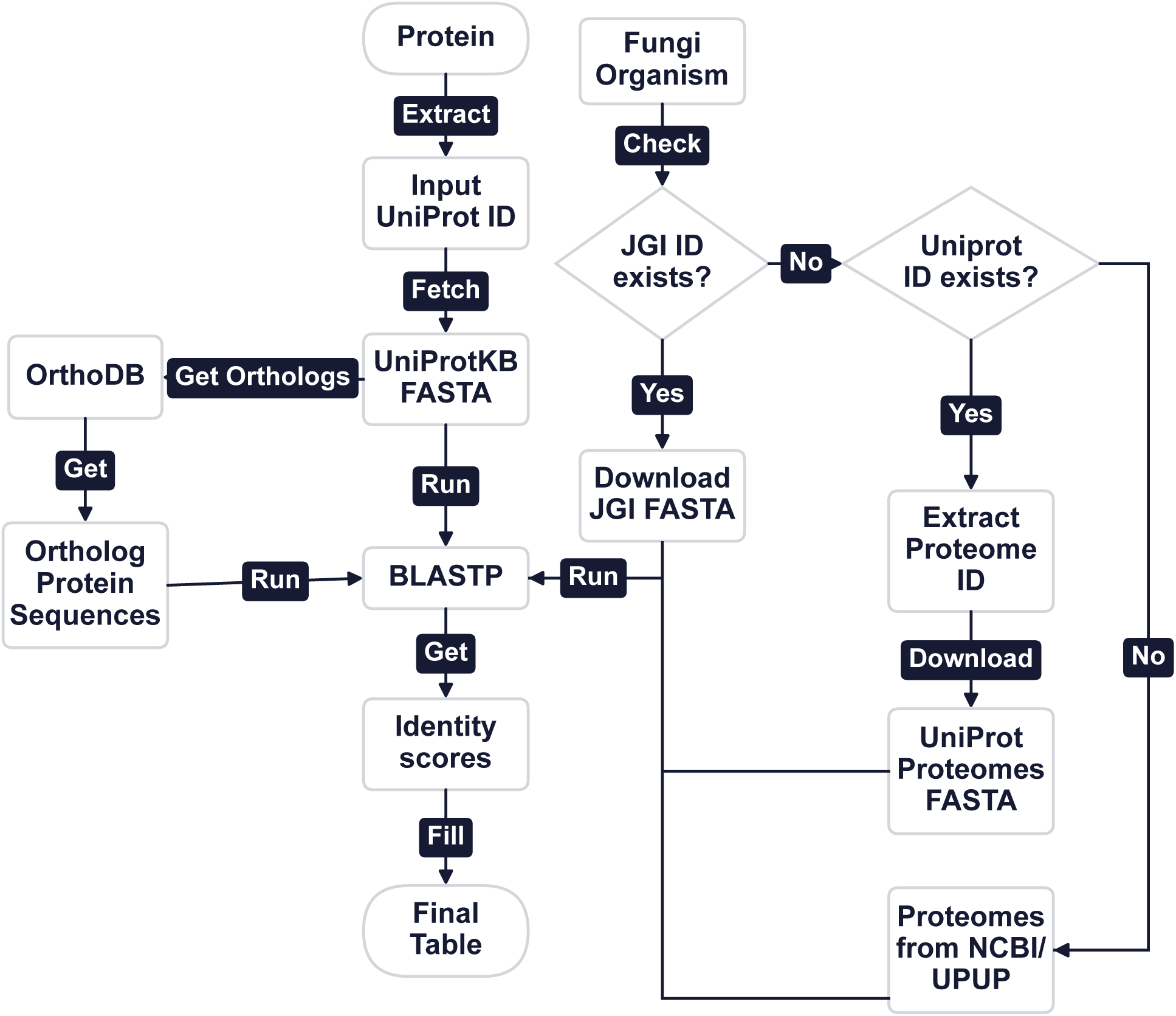
Workflow to create the dataset which describes the contamination potential of various Fungi using the list of 25 proteins in Table 2. Fasta files are downloaded for each of the proteins using their UniProt IDs to then run BLASTp against the proteomes of different Fungi acquired through the different datasources e.g. JGI Genome portal (Nordberg et al., 2014), UniProt Proteomes, OrthoDB tool (Zdobnov et al., 2021).

#### 2.2.1 Compilation of Fungal Species

To construct a comprehensive dataset of fungal species with available proteomes, we employed a multi-source, systematic data retrieval strategy. The compilation includes several databases and repositories to ensure broad coverage of fungal proteomes.

We began with 1,509 fungal accessions from the JGI GOLD database and filtered out duplicates and incomplete entries. For each valid species, proteome FASTA files were retrieved through the JGI Genome Portal API. Where proteomes were unavailable, supplementary data were gathered from UniProt Proteomes, OrthoDB, UniParc, and NCBI RefSeq. In this process we added more species from UniProt that were not in JGI GOLD. This process resulted in a nonredundant dataset of 1,553 unique fungal species with associated proteomes (Figure 2), providing a wide coverage of Ascomycota, Basidiomycota, and other phyla. We reiterate that this process is scalable and extensible and can be used to include additional databases and datasets to grow the list.

**Figure 2:**
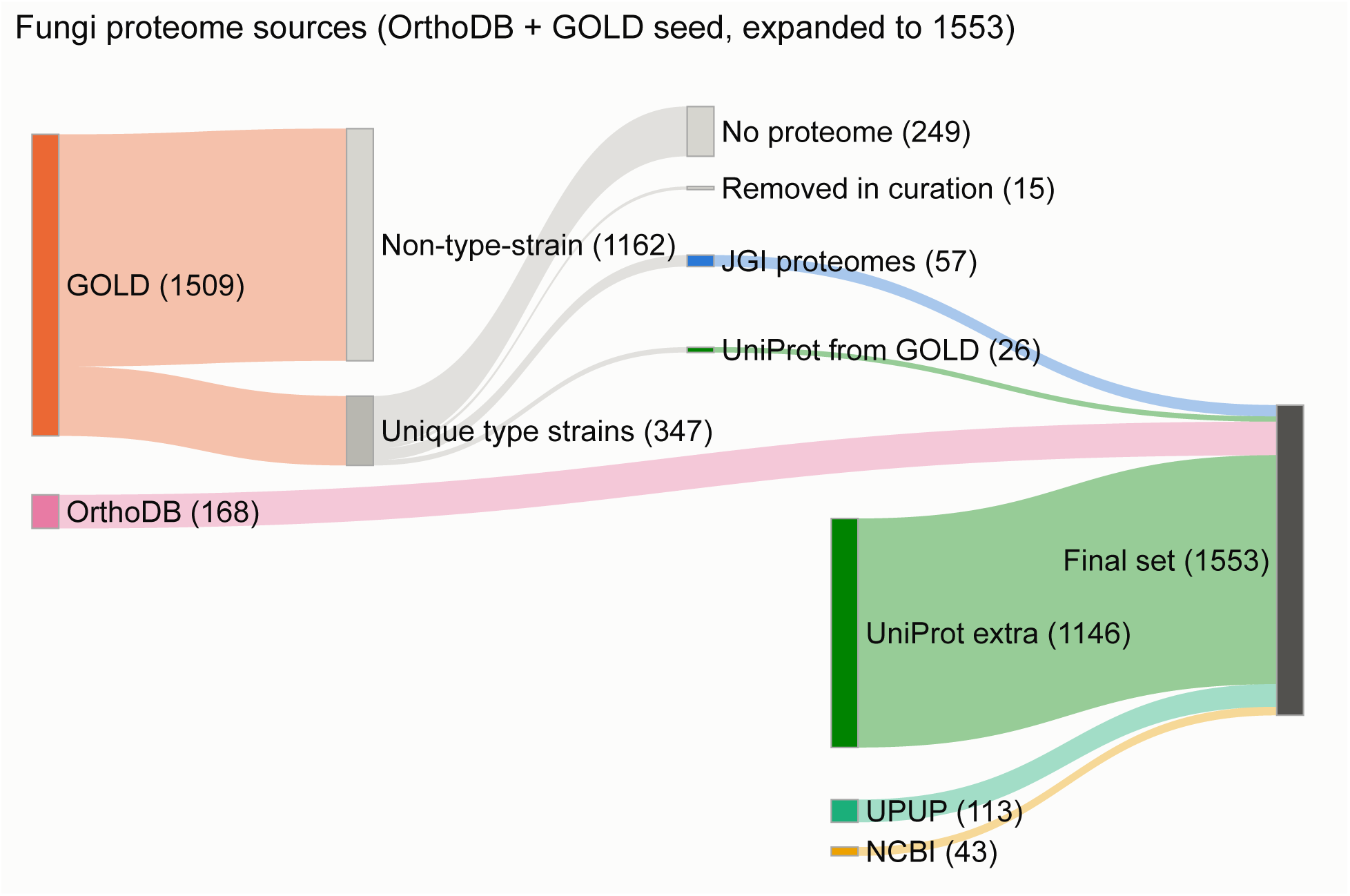
The final fungal dataset in this study was assembled by consolidating entries from several major databases, beginning with 1,509 accessions from the GOLD database that were rigorously filtered to remove duplicates and entries lacking proteome data. Unique GOLD entries and those with available proteomes were supplemented by nonredundant organisms sourced from external repositories, including 1,146 from UniProt, 168 using OrthoDB, 113 from UPUP, and 43 from NCBI. By integrating records across these datasets and ensuring each included organism had proteome-level data, a comprehensive set of 1,553 fungal species were curated.

#### 2.2.2 BLAST Database Creation and Runs

All retrieved proteome FASTA files were used to construct BLAST databases using the NCBI BLAST+ package (Camacho et al., 2009). Individual databases were created per data source to maintain provenance and allow cross-validation of results. Query sequences for the 25 selected proteins were retrieved by UniProtKB identifiers and aligned against each proteome using BLASTp. For each query–subject pair, the maximum sequence identity (*I*) was extracted and used for down-stream scoring. Only the best hit per query protein per organism was retained, ensuring consistency across the dataset.

The identity score (*I*) represents the percentage of identical amino acid residues between the aligned regions of query and subject sequences, normalized by total alignment length (including matches, mismatches, and gaps). Following Peterson et al. (2009), *I ≥* 35% indicates significant structural and functional similarity; this value was adopted as the baseline threshold for ortholog detection in this study as it has survived as an initialization rule for function transfer or SSN exploration. See, for example, Gerlt et al. (2015) and Zallot et al. (2019). At the same time, as described later, we also make use of the more stringent *I ≥* 75% threshold.

When multiple matches were found for the same query protein, the sequence with the highest identity score was selected to represent that pair. It is possible that there are additional protein orthologs with lower *I* that are also indicative of similar traits. Here we do not use them, but generate a table with all orthologs that can be used in future. Some sample orthologs are shown in Table 4. The full list with duplicates removed is also available ^4^.

**Table 4:** A few representative orthologs from our curated set using various query proteins. The *Species* column lists the fungal organism analyzed. The *Q-Protein* column provides the reference protein used for the ortholog search. The new *Q-Organism* column indicates the fungal species origin of the query protein, as parsed from the query protein names. The *Ortholog*column contains the description as present in the header of the proteome FASTA file for the corresponding species. The *Trait* column indicates the property expressed by the query protein (shortforms as in Table 1). Finally, the *I* column shows the percentage sequence identity between the query and the orthologous protein.

| Species | Q-Protein | Ortholog | Q-Organism | Trait | I (%) |
| --- | --- | --- | --- | --- | --- |
| *Candida dubliniensis* | Erg251 | ERG251-like | *Candida albicans* | BF | 94.39 |
| *Cryptococcus gattii* | Plb1 | PLB1-like | *Cryptococcus neoformans* | HP | 91.36 |
| *Metschnikowia* aff. | Efg1 | SOK2 | *Candida albicans* | BF | 89.31 |
| *P. chlamydospora* | Hsp90 | hsc82 | *Aspergillus fumigatus* | TH | 89.09 |
| *Teratosphaeriaceae* sp. | Rad51 | Rhp51 | *Saccharomyces cerevisiae* | RAD | 76.22 |
| *C. cylindrospora* | Cdr1 | AtrD-1 | *Candida albicans* | AMR | 50.47 |
| *Gigaspora rosea* | Lac1 | MCOP | *Cryptococcus neoformans* | HP | 49.46 |
| *Leucoagaricus* sp. | srr1 | TMA-20 | *Coprinopsis cinerea* | SF | 45.67 |
| *Leucoagaricus* sp. | AbaA | TEF-5 | *Emericella nidulans* | SF | 41.53 |

Biopython (Cock et al., 2009) was employed to efficiently handle and parse protein sequences in FASTA format. The *SeqIO* module facilitated reading query and subject sequences, retrieving sequence lengths, and organizing the data for automated downstream processing. This enabled seamless integration of multiple proteomes and streamlined the handling of BLASTp results within the Python workflow.

The results from BLASTp searches across all sources were then combined with previously identified organisms to update the curated list of fungal species. A similar workflow was applied for each data source: proteome extraction, BLASTp alignment, selection of highest-identity matches, and storage of alignments and metadata in a standardized CSV format for downstream analysis.

#### 2.2.3 Scoring

After ortholog identification, the functional overlap of each organism was quantified through two complementary scores: the S-score and A-score, representing breadth and strength of functional similarity, respectively.

Each organism’s contamination potential was summarized using a binary S-score (Equation 1) that indicates the number of functional categories represented by at least one ortholog in that category above the 35% identity threshold. A second metric, the weighted A-score (Equation 2), differentiates between moderate and high sequence similarity by assigning greater weight to properties with *I ≥* 75%. Together, these metrics quantify both the diversity and confidence of stress-associated traits within each proteome.

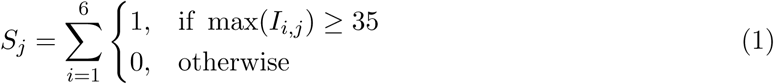

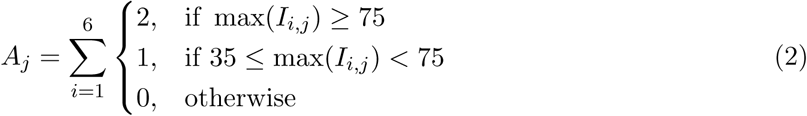

The A-score for an organism ranges from 0 to 12 for the current set of six properties, whereas the S-score ranges from 0 to 6. This is best illustrated by a pair of example species. *Gigaspora margarita* has both S-score and A-score of 6 indicating that while it scores on all six properties considered here, none of them have an identity match above 75%. On the other hand, *Naganishia cerealis* has an S-score of 4 and an A-score of 7 indicating that while it has below 35% identity scores on two of the six properties, three of the four matching properties have a high confidence. Depending on the downstream workflow users can be more aggressive or conservative by choosing one of the two scores, or understanding diversity as well as confidence by using both.

## 3 Results

### Dataset Composition and Taxonomic Coverage

Following integration of datasets from GOLD, UniProt, OrthoDB, UniParc, and NCBI RefSeq, a nonredundant collection of 1,553 fungal species with complete proteomes was obtained (Figure 2). We analyzed orthologs for 25 stress- and virulence-associated proteins across these 1,553 fungal species to evaluate contamination potential in general, but also specifically under spaceflight-relevant conditions. The resulting dataset provides quantitative measures of sequence similarity (S- and A-scores) and trait-level distributions that reveal broad phylogenetic patterns of resilience and risk. This dataset formed the basis for all subsequent analyses of contamination potential.

The taxonomic composition was dominated by Ascomycota (67.2%) and Basidiomycota (21.1%), reflecting the dominance of these phyla in both terrestrial and spacecraft environments. Smaller contributions came from Mucoromycota, Microsporidia, and other early-diverging lineages (Table 5). This wide phylogenetic coverage is in general representative of fungal species studied.

**Table 5:** Distribution of representation percentages and mean A-scores (average ± standard deviation) across fungal phyla.

| Phylum | Counts | Percent counts | A-score (mean ± SD) |
| --- | --- | --- | --- |
| Ascomycota | 1043 | 67.16 | $6.7 \pm 1.4$ |
| Basidiomycota | 327 | 21.06 | $5.7 \pm 1.3$ |
| Mucoromycota | 93 | 5.99 | $5.8 \pm 1.1$ |
| Microsporidia | 31 | 2.00 | $4.2 \pm 1.1$ |
| Glomeromycota | 24 | 1.55 | $6.2 \pm 0.5$ |
| Chytridiomycota | 19 | 1.22 | $6.0 \pm 0.0$ |
| Zoopagomycota | 5 | 0.32 | $5.4 \pm 0.5$ |
| Neocallimastigomycota | 4 | 0.26 | $5.2 \pm 0.5$ |
| Zygomycota | 2 | 0.13 | $4.5 \pm 4.9$ |
| Entomophthoromycota | 2 | 0.13 | $3.0 \pm 4.2$ |
| Blastocladiomycota | 2 | 0.13 | $6.0 \pm 0.0$ |
| Rozellomycota | 1 | 0.06 | $6.0 \pm 0.0$ |

### Scoring Distribution

S-scores were strongly right-skewed (Figure 3, left), with most fungi scoring 5–6, indicating that the majority possessed at least one ortholog above the 35% identity threshold in nearly every functional category. A-scores, which attribute higher weights to high-identity matches, followed an approximately normal distribution centered around 7 (Figure 3, right), showing that moderate sequence conservation is widespread but only a subset of organisms exhibit uniformly high similarity across multiple traits. Together, these patterns reveal that stress-associated functions are broadly distributed in fungi, but few species concentrate multiple high-fidelity adaptations simultaneously.

**Figure 3:**
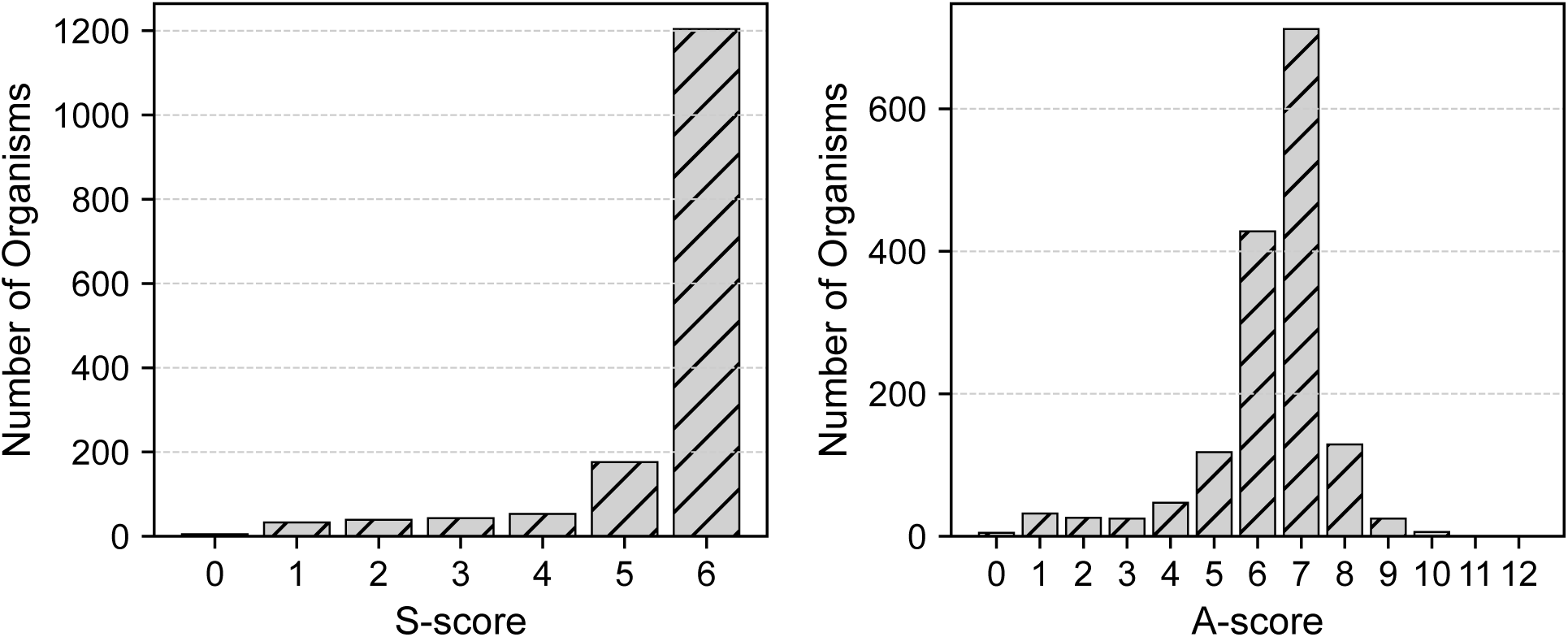
Distribution of S-scores and A-scores for all fungi in our curated dataset of 1,553 fungi. Left: The S-score represents the number of functional categories in which an organism possesses at least one protein with *I ≥* 35. Right: The A-score reflects the weighted sum, giving higher weight to categories with *I ≥* 75, and thus highlights organisms with both broader and higher-confidence functional similarities.

### 3.1 Trait-Level Analysis Across the Dataset

Among the six functional categories, thermophilicity (TH) exhibited the largest number of high-identity matches (*I ≥* 75), encompassing 891 species, far exceeding all other traits (Figure 4, right). Other categories such as biofilm formation (BF), radiation resistance (RAD), and antimicrobial resistance (AMR) showed moderate numbers of high-identity organisms (23–168), whereas most fungi fell within the intermediate identity range (35–75%). This indicates that while thermal resilience is widespread, other stress-tolerance mechanisms are more restricted taxonomically.

**Figure 4:**
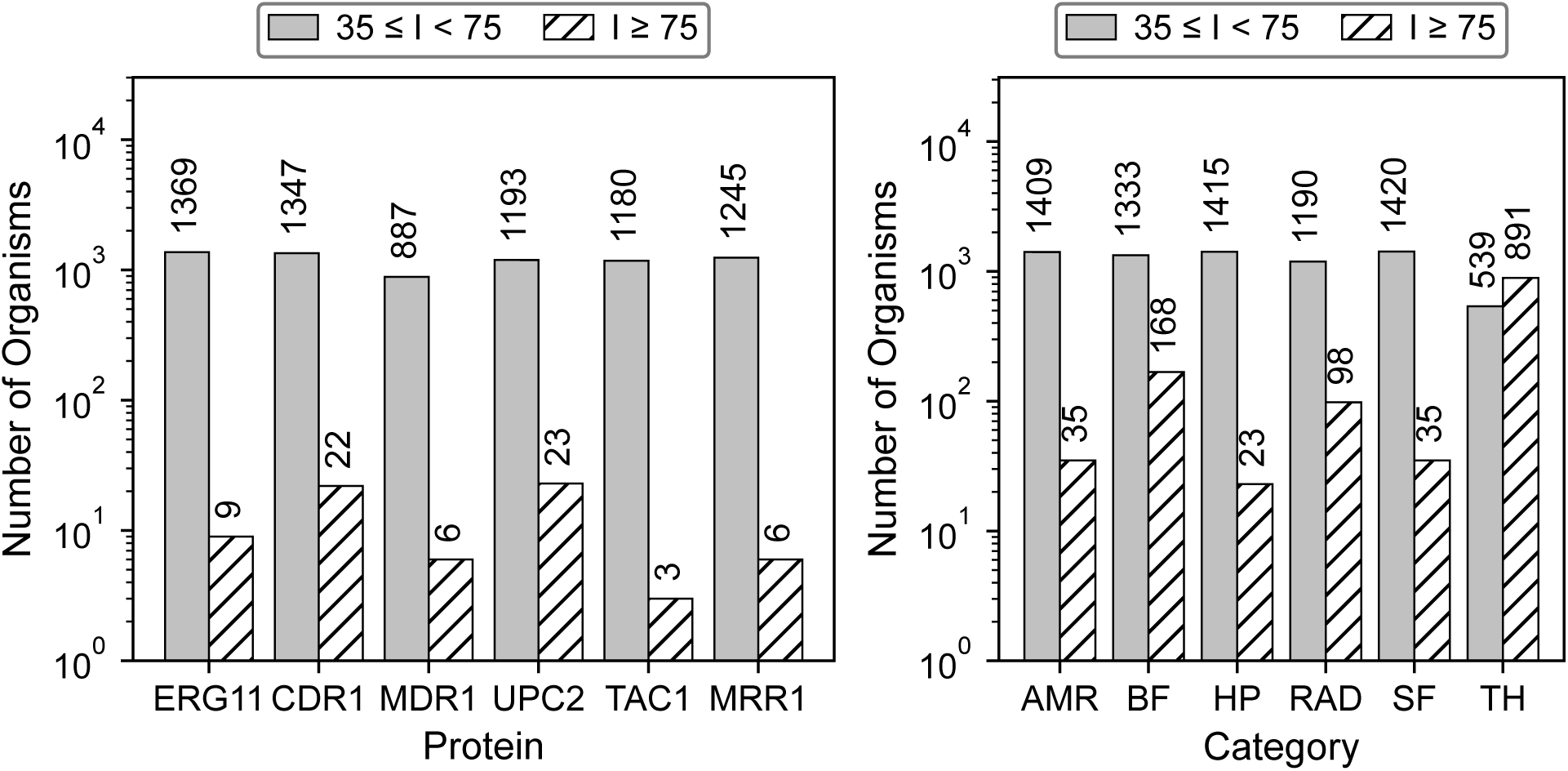
Comparative analysis of identity score distributions for functional protein categories in fungi. Left: Number of unique fungal organisms harboring orthologs to each of six proteins under the Antimicrobial Resistance (AMR) property with *I ≥* 75; a given species may be counted multiple times if it contains high-identity matches to multiple AMR proteins, resulting in a total of 35 unique organisms. Right: Proportion of organisms exhibiting properties known to be potentially harmful, partitioned by high-confidence identity threshold (*I ≥* 75) and moderate-confidence range (35 *≤ I <* 75). Approximately 65.9% fungi have at least one protein with *I ≥* 75 for a harmful property, while about 99.7% reach at least *I ≥* 35, highlighting broad and variable potential for harmful traits within the dataset.

A majority of proteins have orthologs in fewer than 25 organisms in the high-identity group; TAC1, for example, has only 3, and MDR1 only 6. In marked contrast, the medium-index group (35 *≤ I <* 75) consists of several hundred to over a thousand organisms with orthologs per protein/trait, reflecting a wide intermediate as shown in left part of Figure 4.

Trait-level analysis of ortholog presence revealed distinct enrichment patterns (left and right parts of Figure 4). Overall, 99.7% of fungi displayed at least one ortholog with *I ≥* 35%, and 65.9% had at least one with *I ≥* 75%, confirming pervasive potential for stress response traits (Table 6).

**Table 6:** Number of organisms with high (*I ≥* 75) and moderate (35 *≤ I <* 75) sequence identity scores across functional protein categories.

| Category | Moderate ($35 \leq I < 75$) | High ($I \geq 75$) |
| --- | --- | --- |
| Antimicrobial resistance | 1409 | 35 |
| Biofilm formation | 1333 | 168 |
| Human pathogenicity | 1415 | 23 |
| Radiation resistance | 1190 | 98 |
| Spore formation | 1420 | 35 |
| Thermophilicity | 539 | 891 |

These findings indicate that a majority of fungal species harbor at least one functional trait associated with space-relevant environmental resilience.

Across all proteins, only four Ascomycota queries, Efg1, Rim101, Hsp90 and Rad51, showed any *I ≥* 75 hits to organisms outside Ascomycota and Basidiomycota, and these corresponded exclusively to early-diverging fungal lineages. Efg1 and Rim101 each showed a single high-identity ortholog in Mucoromycota (*Rhizopus stolonifer* and *Umbelopsis ramanniana*, respectively). In contrast, Hsp90 from *Aspergillus fumigatus* displayed extensive deep conservation, with 14 matches with *I ≥* 75 spanning Mucoromycota, Glomeromycota, and Zygomycota, while Rad51 from *Saccharomyces cerevisiae* showed three such matches in Zygomycota and Mucoromycota. These protein-specific exceptions align with the broader pattern in Figure 5, where cross-phyla representation largely collapses at the 75% threshold, yet a small set of conserved housekeeping genes retain high-identity orthologs across the deepest branches of the fungal kingdom.

**Figure 5:**
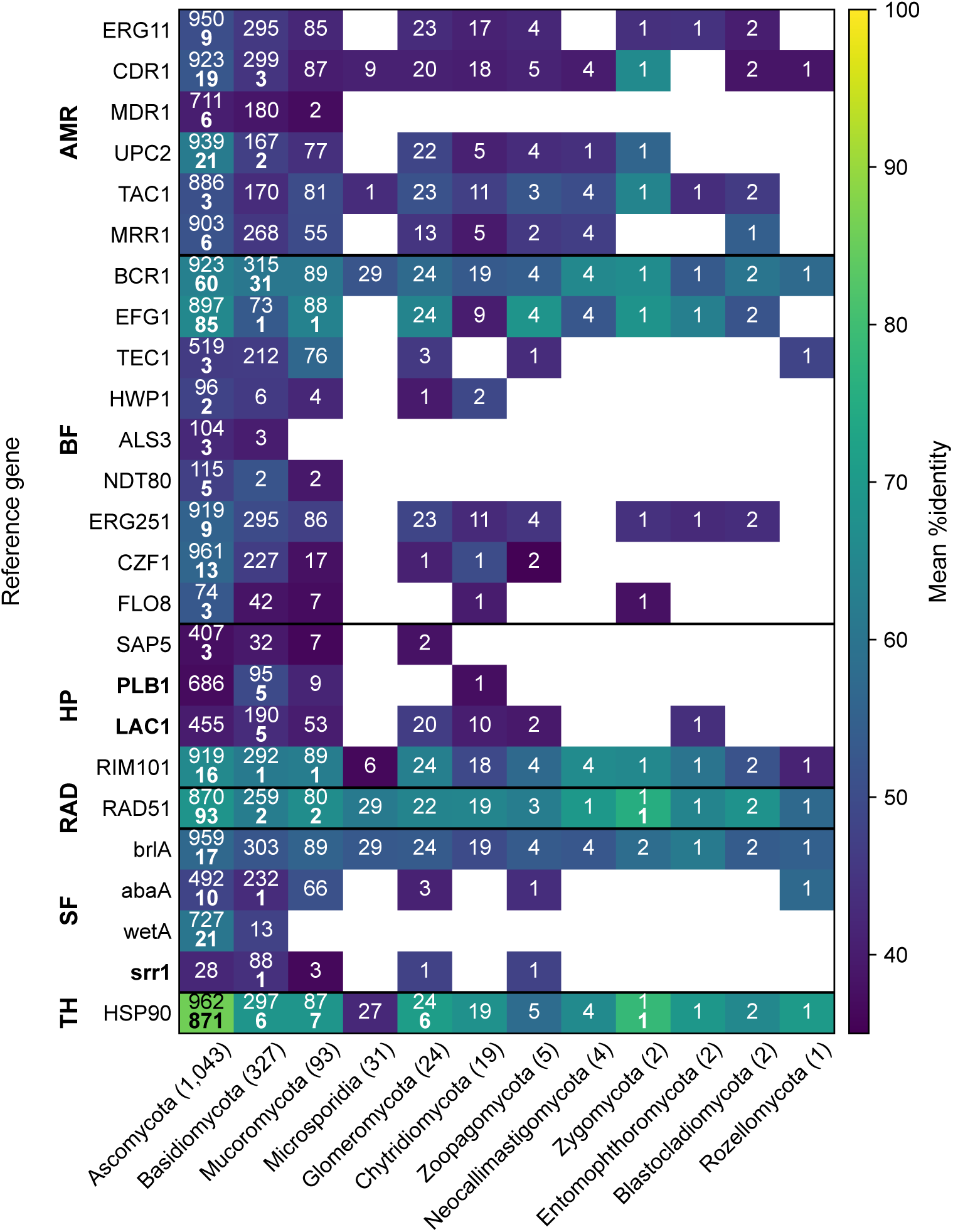
Distribution of BLAST-hit organisms across fungal phyla for each protein at 35% and 75% sequence-identity thresholds. For each phylum the number of curated species is shown immediately after its name (same as in Table 5). For each combination of phylum and reference gene, the corresponding cell shows the number of species that met the two identity thresholds. The color of the cell indicates the mean percentage identity of the species matching the *I ≥* 35% threshold, the mapping shown by the colorbar at the right. Rows are grouped by phenotypic category (AMR, BF, HP, RAD, SF, TH) with horizontal separators between groups; columns are ordered by phylum sample size. The reference for genes shown in bold originates with Basidiomycota species, the other in Ascomycota. The visualization highlights deep-branch retention of HSP90, RAD51, and the spore-development genes (brlA / abaA / wetA), and the near-complete absence of BF and HP orthologs outside Dikarya (Ascomycota and Basidiomycota).

### 3.2 High-Priority Spaceflight Contaminants

When compared with published datasets from the International Space Station and spacecraft assembly facilities (Singh et al., 2018; De Middeleer et al., 2019), several species such as *Penicillium rubens*, *Aspergillus flavus*, and *Rhodotorula* sp. displayed high S and A-scores across multiple functional categories (see Table 7). Although these organisms are BSL-1 and not directly pathogenic, their resilience traits highlight potential for persistent material contamination and interference with spacecraft environmental systems.

**Table 7:** Properties and scores for fungal species found in the Russian Space Station and/or the International Space Station.^a^ The six property columns when added provide A-score given in the first column while their non-zero nature provides the S-score (not separately given). Several of these species have high confidence (*I ≥* 75%) orthologs for thermophilic proteins. We discuss the significance of this in Section 4.

| Species | A-Score | AMR | BF | HP | RAD | SF | TH |
| --- | --- | --- | --- | --- | --- | --- | --- |
| *Alternaria alternata* | 7 | 1 | 1 | 1 | 1 | 1 | 2 |
| *Aspergillus clavatus* | 6 | 0 | 1 | 1 | 1 | 1 | 2 |
| *Aspergillus flavus* | 7 | 1 | 1 | 1 | 1 | 1 | 2 |
| *Aspergillus niger* | 5 | 1 | 0 | 1 | 0 | 1 | 2 |
| *Aspergillus ochraceus* | 7 | 1 | 1 | 1 | 1 | 1 | 2 |
| *Aspergillus versicolor* | 6 | 1 | 0 | 1 | 0 | 2 | 2 |
| *Fusarium* spp. | 6 | 1 | 1 | 1 | 1 | 1 | 1 |
| *Penicillium brevicompactum* | 7 | 1 | 1 | 1 | 1 | 1 | 2 |
| *Penicillium citrinum* | 8 | 1 | 1 | 1 | 1 | 2 | 2 |
| *Penicillium expansum* | 7 | 1 | 1 | 1 | 1 | 1 | 2 |
| *Penicillium nordicum* | 5 | 1 | 1 | 1 | 1 | 1 | 0 |
| *Penicillium rubens* | 7 | 1 | 1 | 1 | 1 | 1 | 2 |
| *Penicillium steckii* | 7 | 1 | 1 | 1 | 1 | 1 | 2 |
| *Rhodotorula* sp. | 6 | 1 | 1 | 1 | 1 | 1 | 1 |
| *Trichosporon asahii* | 2 | 0 | 1 | 1 | 0 | 0 | 0 |

<sup>a</sup> Singh et al.2018, de Midlledeer et al., 2019.

The detection of *P. rubens* and *Rhodotorula* sp. across multiple flight samples and locations, underscores their persistence and adaptability in controlled environments. All detected organisms belonged to BSL-1, but their functional profiles indicate potential operational and material contamination risks under spaceflight conditions.

### 3.3 Network-Level Trait Structure

Given the table of species, phyla, proteins, orthologs etc., a lot of analysis is possible. For instance, a heatmap is shown for Glomeromycota species in Figure 6, where ortholog identity scores for the 25 proteins across the six phenotypic categories is shown for the 24 species in our sample. Species showing putative orthologs to multiple proteins in the same phenotypic category likely represent multiple adaptation strategies. In addition to such heatmaps we make available multiple visualizations on our website where users can also upload their own data to explore patterns. We will keep adding newer plots to the visualizations, including interactive ones like circos.

**Figure 6:**
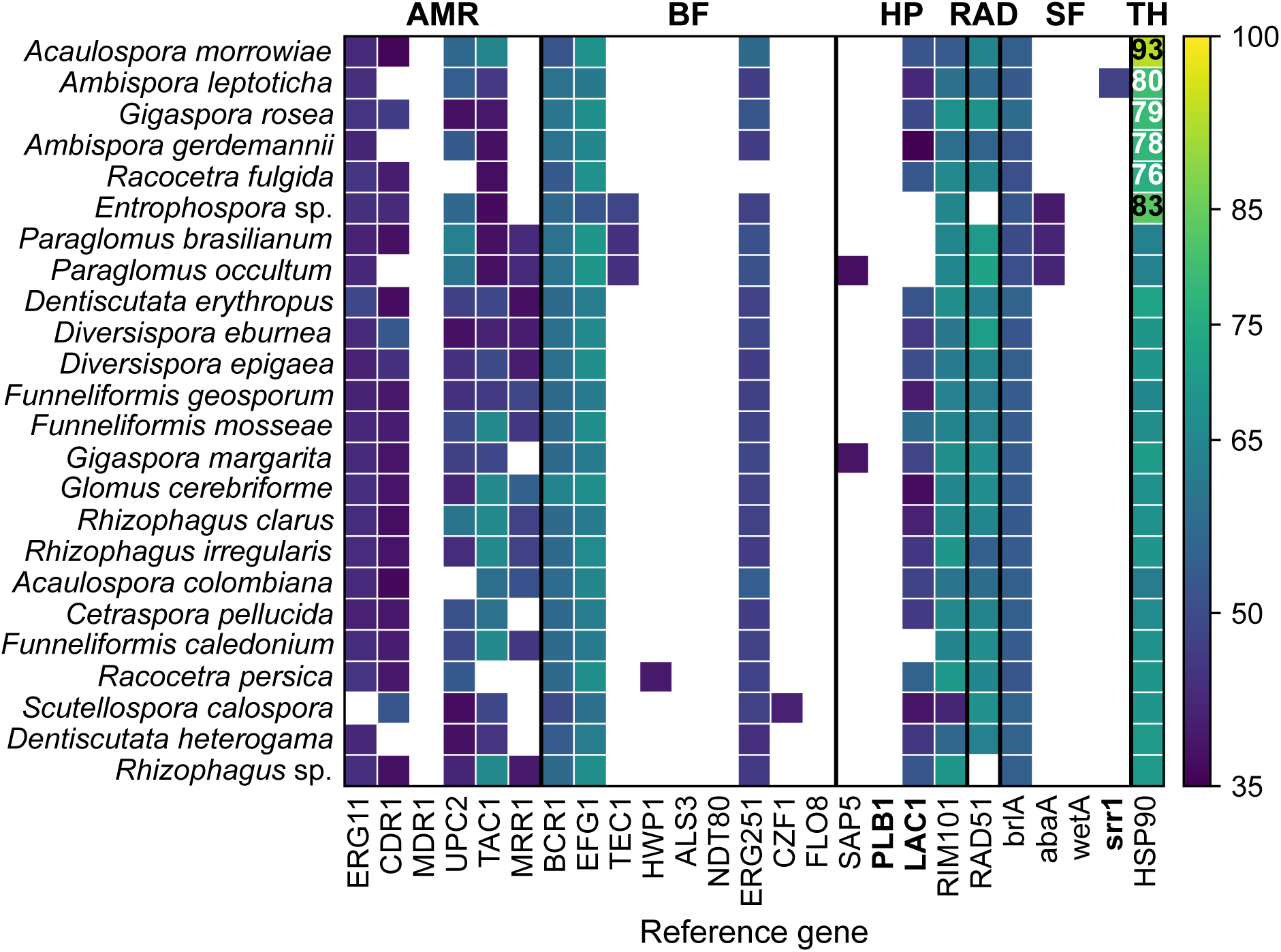
Heatmap of BLASTp percent identity for each of the 24 surveyed Glomeromycota species (rows) against 25 curated reference proteins (columns), drawn from six phenotypic categories: AMR, BF, HP, RAD, SF, and TH considered in this study. Reference proteins drawn from Basidiomycota source organisms are in bold; the remaining 22 are Ascomycota-derived. Cell color encodes %identity; cells with *I <* 35% are left blank. Strong hits (*I ≥* 75%) are annotated with their %identity in bold. Rows are ordered by descending A-score, then by number of strong hits and number of detected orthologs. Across this phylum, strong conservation is confined to HSP90 (6 out of 24). Detection at the looser *I ≥* 35% threshold is broad with 22 of 24 species registering at least one detected ortholog in all six categories. The pattern is consistent with the curated panel’s heavy reliance on Ascomycete-derived references and with the divergent biology of obligate arbuscular mycorrhizal fungi.

## 4 Discussion and Future Directions

While previous microbial monitoring studies have focused primarily on bacteria, fungi represent an underexplored and increasingly relevant concern for planetary protection and astronaut health. The current model of PP bioburden testing done by planetary protection (PP) relies heavily on quantitative culture-based analysis, essentially treating all spore-forming microbial cells as equal liabilities. However, the results of this study’s proteomic trait-based screening suggest that such a "one size fits all" approach is fundamentally incomplete. From a life search or sample-return mission-risk perspective, the threat posed by a contaminant is not merely a function of its abundance, but of its specific biological-trait toolkit. If we consider a science mission to an environment like Mars, characterized by extreme thermal cycling and high ionizing radiation, the biological "cost of entry" for a contaminant is high. A species that is thermophilic but lacks radiation resistance is unlikely to survive transit; on the contrary, a radiation-resistant species that cannot tolerate thermal flux is likely to perish upon landing. The real mission risk, therefore, lies in the overlap of these traits, or the “goldilocks zone of traits”.

When we analyze the results detailed in Table 7 using this perspective, some results are striking. Many species in that list have high confidence orthologs for thermophilic proteins. The emergence of species like *Penicillium citrinum*, *Aspergillus flavus*, and *Rhodotorula* sp. is especially concerning. These are not general contaminants in cleanrooms; they are "biological generalists" with presence across nearly all critical stress categories. The fact that these organisms possess putative orthologs for radiation resistance, thermophilicity, and spore formation at the same time, makes them "perfect candidates" for forward/backward contamination. A spore-forming fungus may survive the vacuum of space and the thermal volatility of a Martian surface will pose the greatest possible risk to the integrity of a landing site, possibly compromising the search for native life by introducing an Earth-based organism capable of persistence. Of the organisms in our curated dataset that have high confidence orthologs for radiation resistance, thermophilicity, none are high confidence spore forming, but there are three viz. *Candida parapsilosis*, *Candida theae*, and *Meyerozyma guilliermondii* that also have high confidence orthologs for biofilm formation, and antimicrobial resistance, and putative orthologs for human pathogenicity and spore formation as well. Besides these 3, there are 38 other species that have high confidence orthologs for radiation resistance and thermophilicity, and putative low confidence orthologs for all of antimicrobial resistance, biofilm formation, human pathogenicity, and spore formation.

Integration of this pipeline into the NASA - PP framework enables future science and crewed missions to shift from a purely descriptive model of cleanliness to a predictive model of cleanliness and risk. This one chance at the basal level converts the Planetary Protection effort from a mere checklist of limited sporeforming biological controls into a precise verification and analysis tool. Instead of just confirming that a spacecraft surface is "clean" based on a culturable spore threshold, we can now assess whether the surviving population has the specific signature needed to survive on the target celestial body. By focusing on this "risk-informed bioburden," we can prioritize elimination of high score taxa, ensuring that the biological signatures we detect are not earth natives.

Our framework offers a first-pass, data-driven screen, but future efforts should aim to couple these computational predictions with functional and environmental validation experiments.

The identified species can have independent uses on Earth through curation processes that lead to bioenergy, biogeochemistry, and bioproducts areas that are of interest to DOE. For instance, *Naganishia tulchinskyi* is capable of forming titan cells (Bijlani et al., 2022) as a stress-response mechanism. This leads to enhanced lipid production, with applications in biofuel generation. Similarly, enzymes from *Aspergillus* species can lead to biomass deconstruction under stress.

Our methodology can be further extended to integrate machine learning models that combine metadata, habitat, and expression profiles to predict contamination risks for previously uncharacterized fungi. Beyond spaceflight, the global rise in temperatures and increasing environmental stressors are likely to enhance fungal adaptability and virulence. Understanding these mechanisms will not only strengthen planetary protection and mission safety but also contribute to broader preparedness against fungal threats in changing terrestrial ecosystems.

## 5 Conclusion

This study demonstrates a scalable and interpretable approach to identifying fungal species of concern in spaceflight environments. By combining curated protein knowledge with ortholog inference across over 1,500 fungal proteomes, we systematically quantify stress and virulence-associated traits even in poorly annotated species. The S- and A-score framework provides a transparent and extensible method for comparative risk assessment.

We also provide an interactive, browser-based tool, CheckFungalContam^5^, that lets users upload species read-counts across one or more locations and returns the per-location contamination potential using thresholds they choose. The tool matches each input against our curated set, reconciles alternate names and duplicates, and presents results as a scored, sortable table, a species-by-location read heatmap, per-property read breakdowns, and an UpSet plot showing which risk properties cooccur. It runs entirely in the browser with no installation. Building on a similar tool originally developed for bacteria (Singh et al., 2022), this establishes a foundation for systematic monitoring and assessment of fungal contamination potential in spaceflight and other extreme environments.

In general, our results aid NASA in defining the next steps in developing a nucleic acid-based spacecraft biological cleanliness verification approach, and understanding host-microorganism risk potentials based on data collected from various spacecraft for PP. The outcomes include defining the technology roadmap of nucleic acid-based technologies, providing an order of magnitude estimate for resource planning to develop this for spacecraft use, developing risk-informed decision-making implementation strategies for human missions, a deeper understanding of standardization on spacecraft surfaces and requirements, and understanding possible gaps in NASA planetary protection toolkit. The tool we have developed will serve as a reference example for other space ferrying agencies like ESA, JAXA, and for private organizations.

## 6 Data and Code Availability

The curated dataset — organism/protein identity matrices, proteome FASTA files, and the synonyms.csv mapping used to consolidate duplicate entries — is archived at CaltechDATA under DOI 10.22002/rttgr-mmc07 (https://doi.org/10.22002/rttgr-mmc07) (concept DOI; resolves to the latest version). The analysis pipeline (proteome retrieval, BLASTp orthology, scoring, and figure generation) is available at https://github.com/FungalMenace/Predicting-Fungal-Contaminants-for-Space-Missions (release v1.0.0; archived DOI: 10.22002/mb4c6-91p24). The interactive tool is available at https://github.com/FungalMenace/CheckFungalContam (release v1.0.0; archived DOI: 10.22002/dzb8t-pyg52) and hosted at https://sites.astro.caltech.edu/checkfungalcontam/. Code is released under the MIT License; the dataset under CC BY 4.0.

## 7 Abbreviations

AI: Artificial Intelligence
AMR: Antimicrobial Resistance
API: Application Programming Interface
BF: Biofilm Formation
Biopython: Bioinformatics Toolkit for Python
BLAST: Basic Local Alignment Search Tool
BLASTp: Basic Local Alignment Search Tool for Proteins
BSL: Bio Safety Level
*C. cylindrospora*: *Coleophoma cylindrospora*
CLI: Command Line Interface
DOE: Department of Energy
ERG251-like: C-4 methyl sterol oxidase
FASTA: Text-based format for representing nucleotide or protein sequences
HP: Human Pathogenicity
HSC82: ATP-dependent molecular chaperone
hsc82 I: Identity Score
ISS: International Space Station
JGI: Joint Genome Institute
MCOP: Multicopper oxidase protein
NASA: National Aeronautics and Space Administration
NCBI: National Center for Biotechnology Information
NID: NASA Interim Directive
NIH: National Institutes of Health
*P. chlamydospora*: *Phaeomoniella chlamydospora*
PLB1-like: Lysophospholipase
RAD: Radiation Resistance
RefSeq: Reference Sequence Database
REST: Representational State Transfer
SF: Spore Formation
TEF-5: Transcriptional enhancer factor - 5
TH: Thermophilicity
TMA: Translation machinery-associated protein

## Acknowledgments

This work was supported in part by the NASA-funded consortium Translational Research Institute for Space Health (TRISH), through the Caltech Space Health Innovation Fund (CSIF). AM and SGD also acknowledge generous support from the Ajax Foundation. We thank the developers of OrthoDB, UniProt, JGI, NCBI and Biopython for maintaining accessible and well-documented APIs. Figures 5 and 6 have been generated with the help of Claude Code using the data that the authors put together. Code for the online app was also generated using Claude Code. We thank the U.S. Department of Energy Joint Genome Institute (JGI) MycoCosm and the principal investigators who generated the unpublished fungal genomes used in this study – Scott Baker, Marcel Bucher, Tadashi Fukami, Chris Todd Hittinger, Li-Jun Ma, Joseph Spatafora, and Huimin Zhao – for permission to use their data. The JGI MycoCosm genomes analyzed and their portals are listed in Supplementary Table 8. The genomics work conducted by the U.S. Department of Energy Joint Genome Institute, a DOE Office of Science User Facility, is supported by the Office of Science of the U.S. Department of Energy under Contract No. DE-AC02-05CH11231.

**Table 8:**
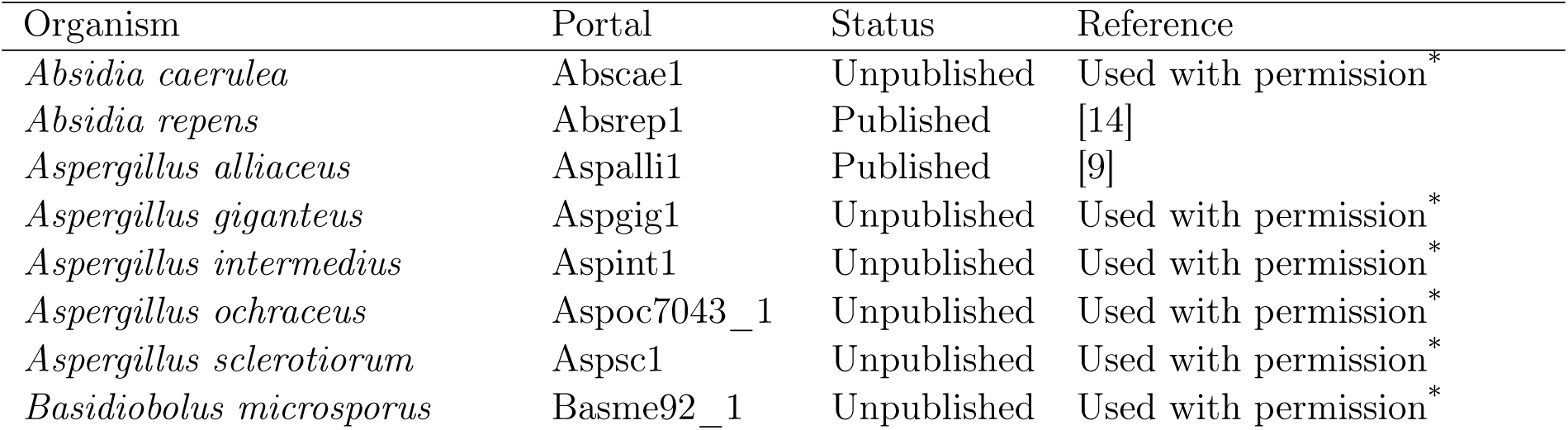
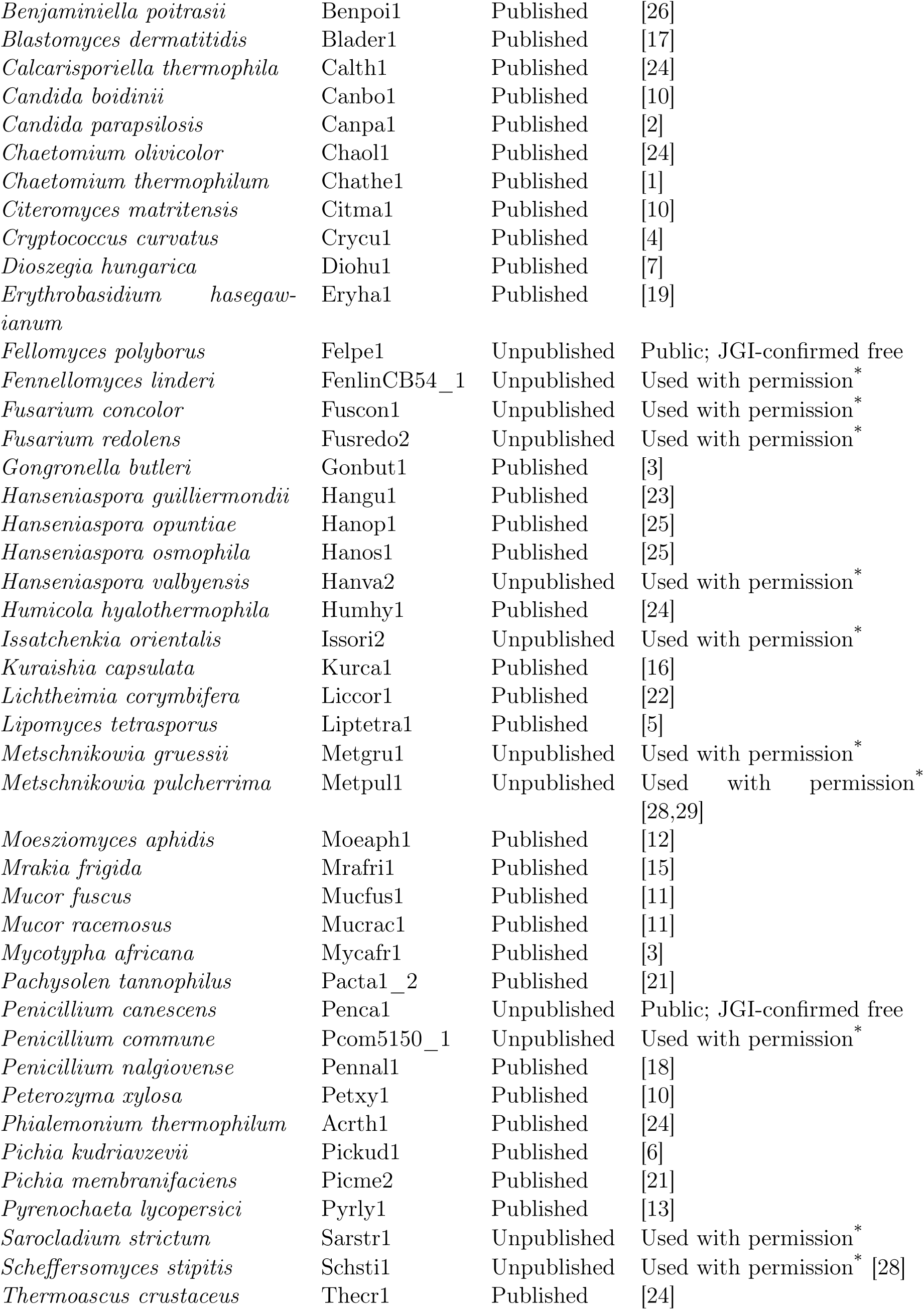
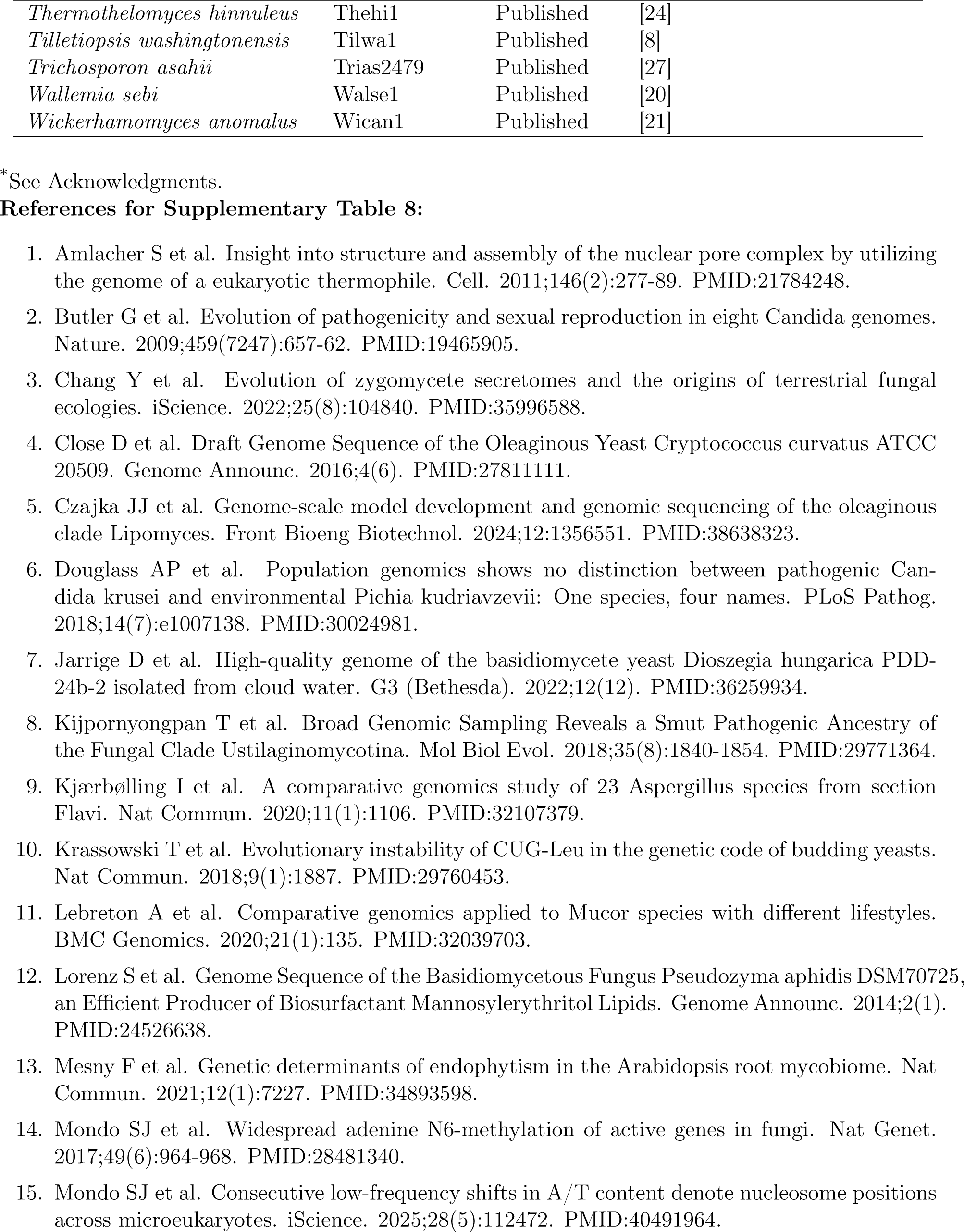
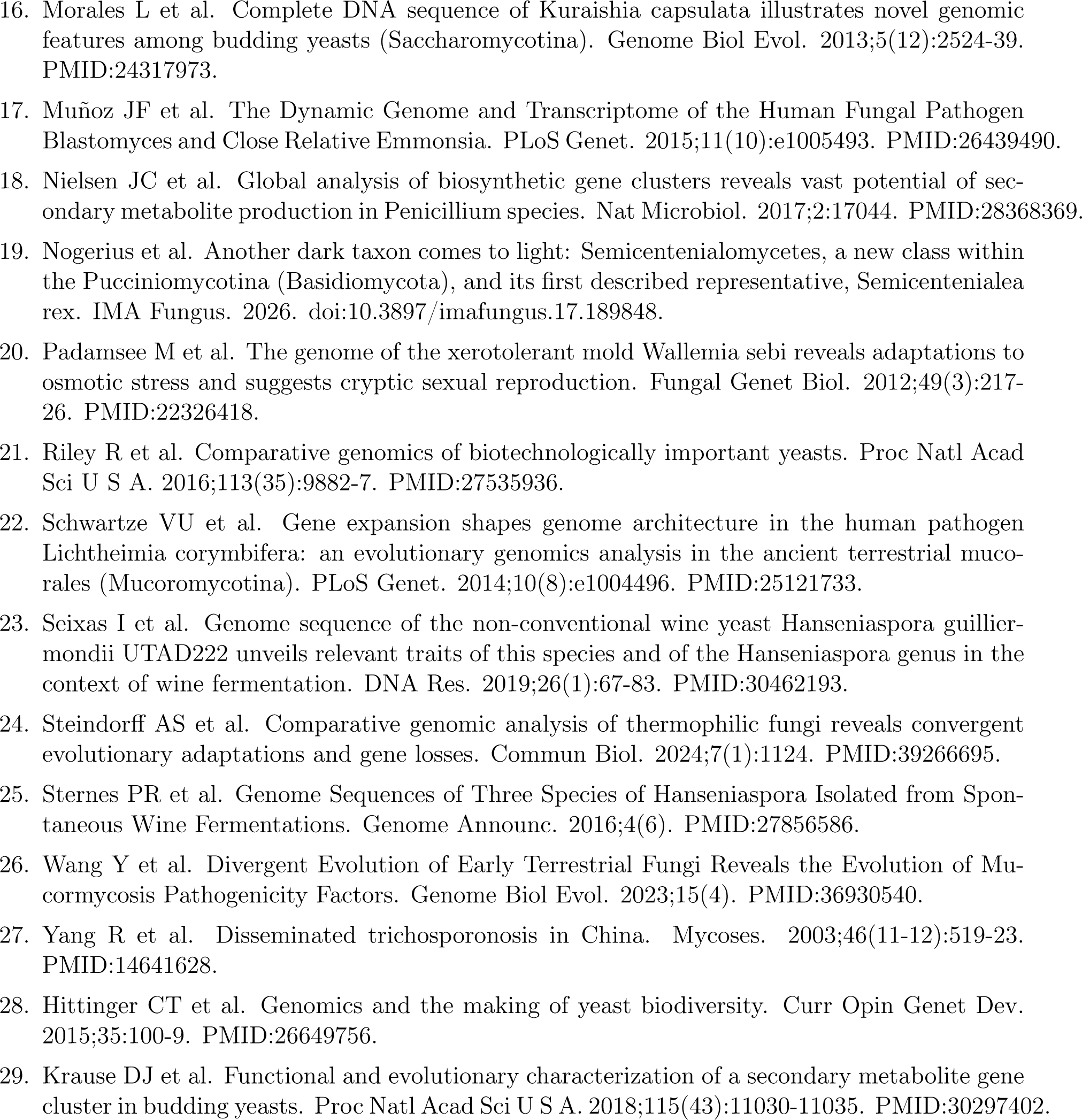
JGI MycoCosm fungal genomes used in this study, referenced by MycoCosm portal (linked in the data deposit, not redistributed). Published genomes cite their primary reference (numbered list below); unpublished genomes were used with permission of the project PIs (see Acknowledgments), except two that JGI confirmed are free to use.

## Footnotes

1 https://www.ncbi.nlm.nih.gov/

2 https://www.uniprot.org/

3 https://humanresearchroadmap.nasa.gov/Gaps/gap.aspx?i=662

4 fungalOrthologs35_1553.csv - part of the data deposit at https://doi.org/10.22002/rttgr-mmc07.

5 https://sites.astro.caltech.edu/checkfungalcontam/ (source: https://github.com/FungalMenace/CheckFungalContam).

## Notes

https://doi.org/10.22002/rttgr-mmc07

https://github.com/FungalMenace/Predicting-Fungal-Contaminants-for-Space-Missions

https://sites.astro.caltech.edu/checkfungalcontam/

https://github.com/FungalMenace/CheckFungalContam

### Competing Interest Statement

The authors have declared no competing interest.

